# Consolidation and maintenance of *Drosophila* long-term memory require LIM homeodomain protein Apterous in distinct brain neurons

**DOI:** 10.1101/2020.09.29.319715

**Authors:** Show Inami, Tomohito Sato, Yuki Suzuki, Toshihiro Kitamoto, Takaomi Sakai

## Abstract

The LIM-homeodomain (LIM-HD) transcription factor Apterous (Ap) and its cofactor Chip (Chi) form a complex that regulates various developmental events in *Drosophila*. Although Ap continues to be expressed in the adult brain, the functions of the centrally expressed Ap remain incompletely understood. Here, we show that Ap and Chi in the *Drosophila* memory center, the mushroom bodies (MBs), are indispensable for long-term memory (LTM) maintenance, whereas Ap in a subset of clock neurons [large ventral-lateral neurons (l-LNvs)] plays a crucial role in memory consolidation in a Chi-independent manner. *Ex vivo* imaging revealed that Ap, but not Chi, in l-LNvs is essential for the appropriate Cl^−^ responses to GABA. Furthermore, knockdown of GABA_A_ receptor in l-LNvs compensated for the impairment of memory consolidation in *ap* null mutant flies. Our results indicate that *Drosophila* Ap functions differently in l-LNvs and MBs, and it contributes to the consolidation and maintenance of LTM.

## Introduction

*Drosophila* Apterous (Ap) is one of the most well-studied LIM-homeodomain (LIM-HD) proteins. Similarly to other LIM-HD proteins, Ap has two protein–protein interaction domains (LIM) and a DNA-binding homeodomain motif (HD). Ap coordinates with its cofactor Chip (Chi), and the multimeric Ap/Chi complex (hereafter referred to as Ap/Chi) acts as a positive regulator of transcription (Hobert and Westphal, 2000). In the fruitfly *Drosophila melanogaster*, *ap* was initially identified as a regulatory gene in wing development (Cohen et al., 1992; Diazbenjumea and Cohen, 1993). The subsequent investigations revealed that Ap and Chi are also essential for nervous system development (Lundgren et al., 1995; O’Keefe et al., 1998; Hobert and Westphal, 2000; van Meyel et al., 2000; Stratmann et al., 2019). Similarly, Lhx2, a mammalian ortholog of Ap, plays essential roles in neurodevelopmental events (e.g., cell proliferation, axon pathfinding, and neurite outgrowth) in the central nervous system (Rincon-Limas et al., 1999; Hobert and Westphal, 2000; Chou and Tole, 2019). In particular, Lhx2 is involved in the development of the mouse hippocampus, which is one of the crucial brain structures regulating learning and memory (Sanuki et al., 2011; Subramanian et al., 2011; Abellan et al., 2014; Godbole et al., 2018; Chou and Tole, 2019). Lhx2 continues to be expressed in the mature hippocampus (Lakhina et al., 2013), indicating its role in hippocampus-dependent brain functions such as learning and memory. However, little is known about Lhx2 functions in the adult hippocampus. As was observed in mammalian Lhx2, Ap also continues to be expressed in many neurons in the adult fly brain. Although it has been shown that Ap in the adult brain neurons expressing a neuropeptide pigment-dispersing factor (Pdf) is required for proper sleep/wake regulation, functions of the centrally expressed Ap remain largely unclarified (Shimada et al., 2016).

In this study, using a courtship conditioning assay, we examined if and how Ap in the adult brain is involved in *Drosophila* learning and memory. In the courtship conditioning assay, male flies are paired with unreceptive mated females, which give sufficient stresses (e.g., courtship-inhibiting cues and sexual rejection) to males to interfere with mating success (conditioning). After being conditioned, their memory is subsequently observed as suppression of male courtship even toward virgin females. A newly formed labile memory is consolidated into a more stable long-term memory (LTM) in the brain. In courtship conditioning, several genes involved in memory consolidation to establish LTM have been identified. These include (1) *Cyclic-AMP response element-binding protein B* (*CrebB*) encoding a transcriptional factor (Sakai et al., 2004), (2) *Notch* encoding a transmembrane receptor (Presente et al., 2004), (3) *orb2* encoding an mRNA binding protein (Keleman et al., 2007), and (4) *Ecdysone Receptor* encoding a nuclear hormone receptor (Ishimoto et al., 2009). Since these gene products regulate protein expression, it is most likely that *de novo* protein synthesis is essential for memory consolidation to establish LTM induced by courtship conditioning (hereafter, referred to as courtship LTM). Once consolidated, LTM requires continual maintenance until recall, because memory traces gradually decay via molecular turnover and cellular reorganization. In aversive olfactory memory or courtship memory, LTM maintenance requires transcriptional activation of CREB in the *Drosophila* memory center, the mushroom bodies (MBs) (Hirano et al., 2016; Inami et al., 2020), suggesting that novel protein synthesis via transcription is also necessary for the LTM maintenance. However, little is known about the genetic and cellular underpinning regulating LTM maintenance.

Here, we provide the evidence that Ap and Chi in the MBs are essential for the maintenance of courtship LTM, whereas Ap in Pdf neurons is necessary for memory consolidation through the regulation of the appropriate Cl^−^ responses to GABA in a Chi-independent manner.

## Results

### *ap* mutants are defective in LTM

We examined whether Ap is involved in *Drosophila* memory using a courtship conditioning assay. In this assay, 1 h conditioning generates short-term memory (STM), and 7 h conditioning induces LTM (Siegel and Hall, 1979; Sakai et al., 2004; Sakai et al., 2012). STM persists for at least 8 h, whereas LTM persists for at least 5 d (Sakai et al., 2004; Sakai et al., 2012). Two mutant alleles of *ap* (*ap^rk568^* and *ap^UGO35^*) were tested for the effects of *ap* mutations on LTM. *ap^rk568^* is a strong hypomorphic allele with a P element insertion into the putative 5’-flanking region (Cohen et al., 1992). *ap^UGO35^* (hereafter referred to as *ap^null^*) is a null allele, generated by imprecise excision of the P element using *ap^rk568^* (Cohen et al., 1992). Since *ap^rk568^* and *ap^null^* are homozygous lethal, we used heterozygous mutants (*ap^rk568^*/+ and *ap^null^*/+). In wild-type, *ap^rk568^*/+, and *ap^null^*/+ flies, the courtship index (CI), which is an indicator of male courtship activity, of conditioned males was significantly lower than that of naïve males 1 h after 1 h conditioning (Fig. 1*A*, CI). To quantify courtship memory, memory index (MI) was calculated (refer to Materials and Methods). In MI, no significant differences were detected among three genotypes (Fig. 1*A*, MI; WT vs *ap^rk568^*/+, Permutation test; *P* = 0.550; WT vs *ap^null^*/+, Permutation test; *P* = 0.896). Thus, in *ap^rk568^*/+ and *ap^null^*/+, STM was intact. On day 5 after 7 h conditioning (hereafter referred to as 5 d memory), wild-type flies also showed a significant reduction in CI in a conditioning-dependent manner (Fig. 1*B*, CI). However, no significant differences in CI were detected between naïve and conditioned *ap^rk568^*/+ and *ap^null^*/+ (Fig. 1*B*, CI), and their LTM was impaired (Fig. 1*B*, MI; WT vs *ap^rk568^*/+, Permutation test; *P* = 0.004; WT vs *ap^null^*/+, Permutation test; *P* = 0.015), indicating that Ap is essential for LTM.

**Figure 1.**
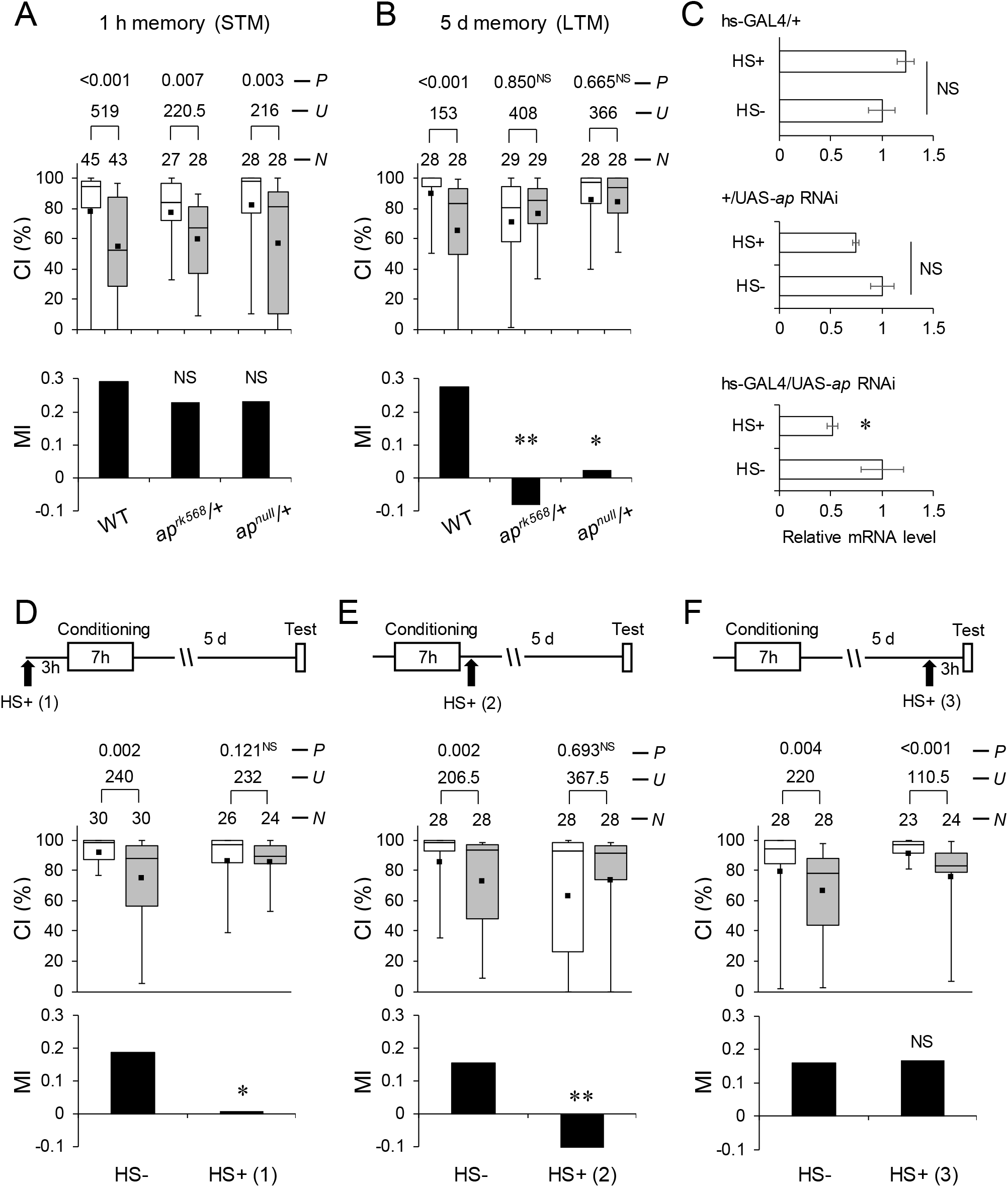
Ap is necessary for LTM. ***A***, Wild-type, *ap^rk568^*/+, and *ap^null^*/+ flies were used in the experiments. Males were tested for 1 h after 1 h conditioning (1 h memory). ***B***, Wild-type, *ap^rk568^*/+, and *ap^null^*/+ flies were used in the experiments. Males were tested on d 5 after 7 h conditioning (5 d memory). ***C***, Real-time qRT-PCR analysis of *ap* mRNA expression level. hs-GAL4 was used for the induction of *ap* RNAi. Mean ± SEM was calculated from five to six replicates. HS-, non-heat-shocked flies. HS+, flies with heat-shock treatment (20 min) 3 h before RNA extraction. *, *P* < 0.05; NS, not significant. ***D***, 5 d memory after 7 h conditioning in flies with or without a 20 min heat-shock treatment 3 h before conditioning [HS+ (1)]. ***E***, 5 d memory after 7 h conditioning. HS+ (2), flies with heat-shock treatment immediately after conditioning. ***F***, 5 d memory after 7 h conditioning. HS+ (3), flies with heat-shock treatment 3 h before test. ***A***, ***B***, **and *D***–***F***, Box plots for a set of CI data show 5th, 25th, 75th, and 95th centiles. The white boxes indicate naïve males and the gray boxes indicate conditioned males. CI, courtship index; MI, memory index; *N*, sample size; *U*, Mann– Whitney *U*; *P*, probability. *, *P* < 0.05; **, *P* < 0.01; NS, not significant. ***D***–***F***, hs-GAL4/UAS-*ap* RNAi flies were used.

### Ap expression during the adult stage is necessary for LTM

Since Ap plays essential roles in neurodevelopmental events in the central nervous system (Rincon-Limas et al., 1999; Hobert and Westphal, 2000; Chou and Tole, 2019), it is possible that the observed LTM impairment is caused by neurodevelopmental defects in *ap* mutant flies. To determine if the *ap* expression in adulthood is critical for LTM, we examined the effects of temporal knockdown of *ap* expression on the LTM phenotype using flies heterozygous for heat-shock (hs)-GAL4 and UAS-*ap* RNAi lines (hs-GAL4/UAS-*ap* RNAi). First, the effectiveness of *ap* RNAi was confirmed by qRT-PCR. Flies were heat-shocked (37 °C) for 20 min, and total RNA was extracted 3 h after heat-shock treatment. Heat-shocked hs-GAL4/UAS-*ap* RNAi flies showed about 50% reduction in *ap* expression level compared with control flies (Fig. 1*C*; hs-GAL4/+, Student’s *t*-test, *t*_(8)_ = −1.474, *P* = 2.306; +/UAS-*ap* RNAi, *U* = 5, *P* = 0.117; hs-GAL4/UAS-*ap* RNAi, *U* = 4, *P* = 0.025). In the absence of heat-shock treatment, LTM in hs-GAL4/UAS-*ap* RNAi flies was detected [Fig. 1*D–F*; CI in HS-(1), *U* = 240, *P* = 0.002; CI in HS-(2), *U* = 206.5, *P* = 0.002; CI in HS-(3), *U* = 220, *P* = 0.004]. In contrast, LTM impairment was observed when flies were heat-shocked 3 h before or immediately after 7 h conditioning (Fig. 1*D, E*; HS−(1) vs HS+(1), Permutation test; *P* = 0.037; HS−(2) vs HS+(2)., Permutation test; *P* = 0.007). These results demonstrate that normal Ap function is required for the consolidation and maintenance of LTM. However, when flies were heat-shocked 3 h before the test, LTM was found to be intact (Fig. 1*F*; HS−(3) vs HS+(3), Permutation test; *P* = 0.956). On the basis of the results of qRT-PCR analysis (Fig. 1*C*), the *ap* expression level must be reduced to ~50% of the control level during the test (3 h after heat-shock treatment). Therefore, reduced Ap function does not have a critical effect on LTM retrieval. Under this condition, the *ap* expression level may not be reduced to a sufficient degree or duration to disrupt the LTM maintenance phase.

### Ap in MB α/β neurons is indispensable for memory maintenance

To examine the significance of Ap in the adult brain for memory consolidation and/or maintenance, we next employed UAS-*ap* RNAi in combination with three GAL4 lines driving GAL4 expression in different neuronal subsets of MBs, the essential brain structure for courtship LTM (Ishimoto et al., 2009; Sakai et al., 2012; Fitzsimons et al., 2013; Inami et al., 2020). LTM was intact when Ap was knocked down in MB γ or α’/β’ neurons, whereas Ap knockdown in MB α/β neurons impaired LTM (Fig. 2*A*, MI; *R41C10*/+ vs *R41C10* / UAS-*ap* RNAi, Permutation test, *P* = 0.011; UAS-*ap*RNAi/+ vs *R41C10* / UAS-*ap* RNAi, Permutation test; *P* = 0.033).

**Figure 2.**
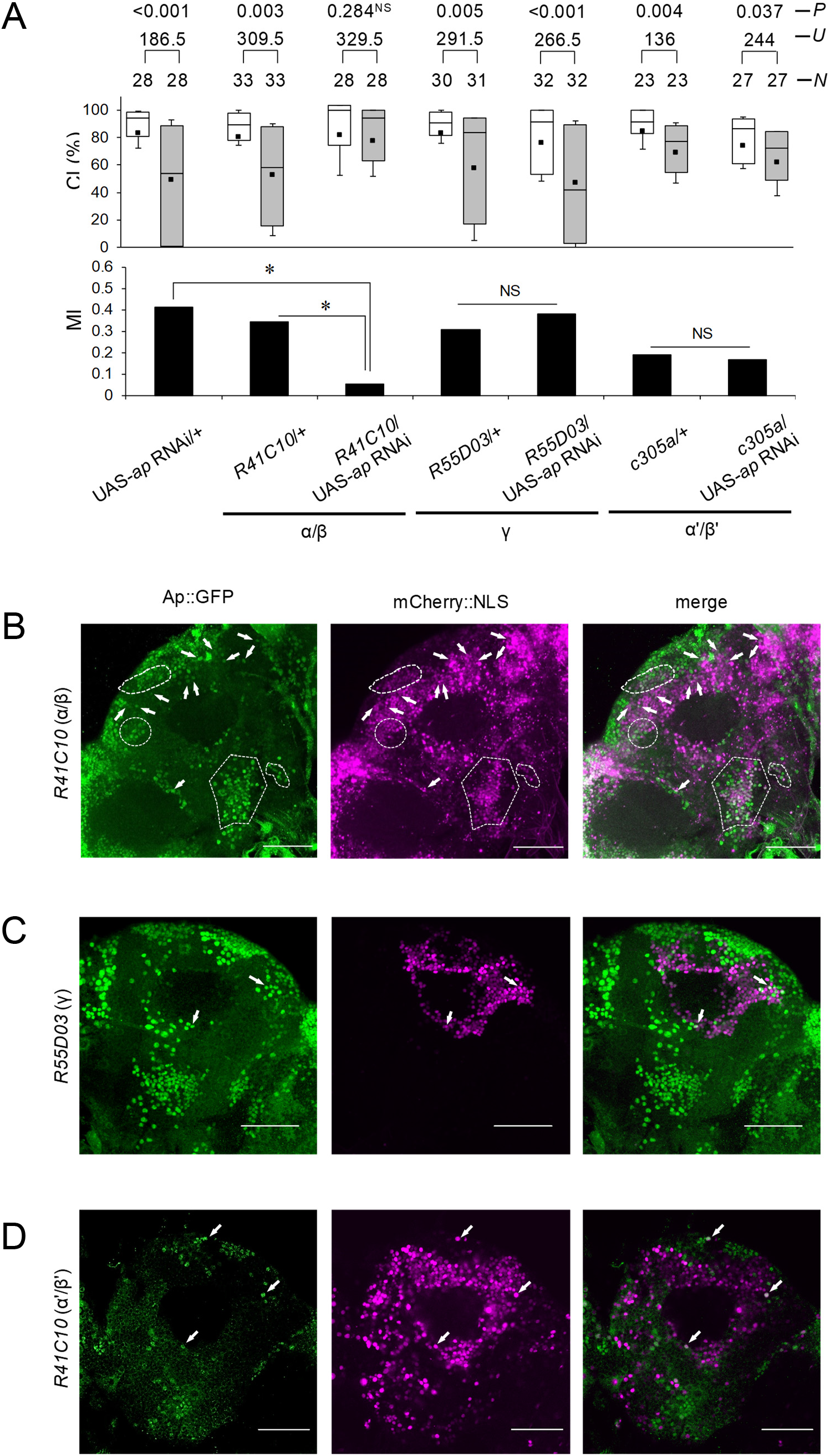
*ap* knockdown in MB α/β neurons induces LTM impairment. ***A***, 5 d memory after 7 h conditioning. *ap* RNAi was driven by MB α/β-GAL4 (*R41C10*), MB γ-GAL4 (*R55D03*), and MB α’/β’-GAL4 (c305a). *, *P* < 0.05; NS, not significant. ***B***–***D***, Ap-expressing neurons in MB neurons. Confocal sectional images at the level of Kenyon cells in MBs. Scale bars, 50 μm; green, Ap::GFP; magenta, mCherry::NLS driven by MB-GAL4 lines. Dotted lines and arrows show colocalization of Ap::GFP and mCherry::LNS. ***B***, *ap::GFP*/+; UAS-*mCherry::NLS*/*R41C10* flies were used. ***C***, *ap::GFP*/*+*; UAS-*mCherry::NLS*/*R55D03* flies were used. ***D***, *ap::GFP*/*c305a*; UAS-*mCherry::NLS*/*+* were used.

We next examined whether Ap-expressing neurons are present in the brain area where somata of the MB neurons (Kenyon cells) are located. To visualize Ap-expressing neurons, we used *ap::GFP* knock-in flies, which express a GFP reporter in a pattern consistent with endogenous Ap expression (Caussinus et al., 2011). Ap::GFP and the nucleus-targeted mCherry reporter for MB α/β neurons were found to be colocalized in many neurons (Fig. 2*B*), and only a few neurons showed colocalization in MB γ or α’/β’ neurons (Fig. 2*C, D*).

To clarify whether Ap in MB α/β neurons is responsible for memory consolidation or maintenance, we performed temporal knockdown of *ap* in MB α/β neurons using the TARGET system (McGuire et al., 2003). To knockdown *ap* during the memory consolidation or maintenance phase, the temperature was increased to 30 °C for 24 h during two different experimental periods: (1) starting at 24 h before the end of conditioning [Fig. 3*A*, RT(1)], and (2) starting at 48 h after the end of conditioning (memory maintenance phase) [Fig. 3*A*, RT(2)]. LTM was impaired when Ap was knocked down during the memory maintenance phase, but it was intact when Ap was knocked down during the conditioning phase [Fig. 3*C*; Permutation test; PT vs RT(1), *P* = 0.3274; PT vs RT(2), *P* < 0.001]. In control flies, the temperature shift did not affect LTM [Fig. 3*B*; Permutation test; PT vs RT(1), *P* = 0.968; PT vs RT(2), *P* = 0.602]. Thus, Ap in MB α/β neurons mainly contributes to LTM maintenance rather than memory consolidation.

**Figure 3.**
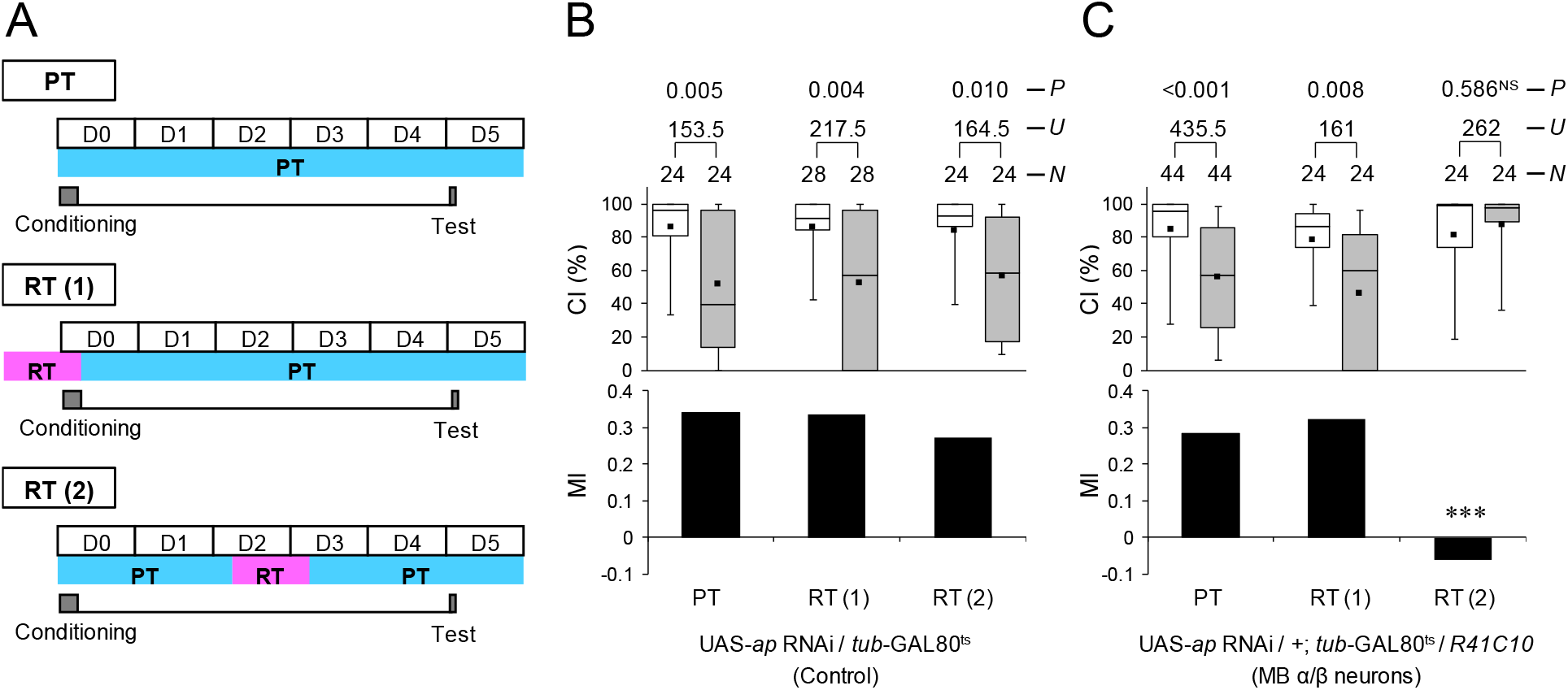
Temporal *ap* knockdown in MB α/β neurons or in Pdf neurons. ***A***, Experimental paradigms for temperature shift treatments. PT, permissive temperatures (25 °C). RT (1), flies were kept at restrictive temperature (RT, 30 °C) for 24 h before the end of conditioning. RT (2), flies were kept at RT for 48–72 h after 7 h conditioning. ***B and C***, 5 d memory after 7 h conditioning. ***, *P* < 0.001; NS, not significant. ***B***, UAS-*ap* RNAi/*tub*-GAL80ts flies were used as a control. ***C***, UAS-*ap* RNAi/+; *tub*-GAL80^ts^/*R41C10* flies were used.

### Ap in Pdf neurons is indispensable for memory consolidation

Ap knockdown by hs-GAL4 shows that Ap is involved in memory consolidation and maintenance (Fig. 1). Nevertheless, Ap in MB α/β neurons mainly contributed to LTM maintenance (Fig. 3*C*). Thus, it is possible that brain neurons other than the MB α/β neurons are involved in Ap-dependent memory consolidation. We previously identified that Ap is expressed in Pdf neurons, which consist of two neural clusters [small ventral–lateral neurons (s-LNvs) and large ventral–lateral neurons (l-LNvs)](Shimada et al., 2016). Thus, we next examined whether *ap* knockdown in Pdf neurons affects LTM. The following three GAL4 lines were used: (1) *Pdf*-GAL4, which drives GAL4 expression in both s-LNvs and l-LNvs, (2) *c929*, which drives GAL4 expression in peptidergic neurons including l-LNvs (Taghert et al., 2001), and (3) *R18F07*, which drives GAL4 expression in s-LNvs and only weakly in one of the l-LNvs (Inami et al., 2020). When *Pdf*-GAL4 and *c929* were used to knock down *ap*, LTM was impaired (Fig. 4*A*; Permutation test; in *Pdf*-GAL4/UAS-*ap* RNAi, vs UAS control, *P* < 0.001, vs GAL4 control, *P* < 0.001; in *c929*/UAS-*ap* RNAi, vs UAS control, *P* < 0.001, vs GAL4 control, *P* = 0.001), whereas *ap* knockdown by *R18F07* did not (Fig. 4*A*; in *R18F07*/UAS-*ap* RNAi, vs UAS control, *P* = 0.099, vs GAL4 control, *P* = 0.916). These results suggest that Ap in l-LNvs is necessary for memory consolidation. Thus, we examined when *ap* in Pdf neurons is required during memory processing for LTM. In contrast to temporal knockdown in the MB α/β neurons, LTM was impaired when Ap was knocked down during conditioning but not during maintenance (Fig. 4*B*; Permutation test; *P* = 0.019). These findings indicate that Ap in Pdf neurons mainly contributes to memory consolidation rather than LTM maintenance.

**Figure 4.**
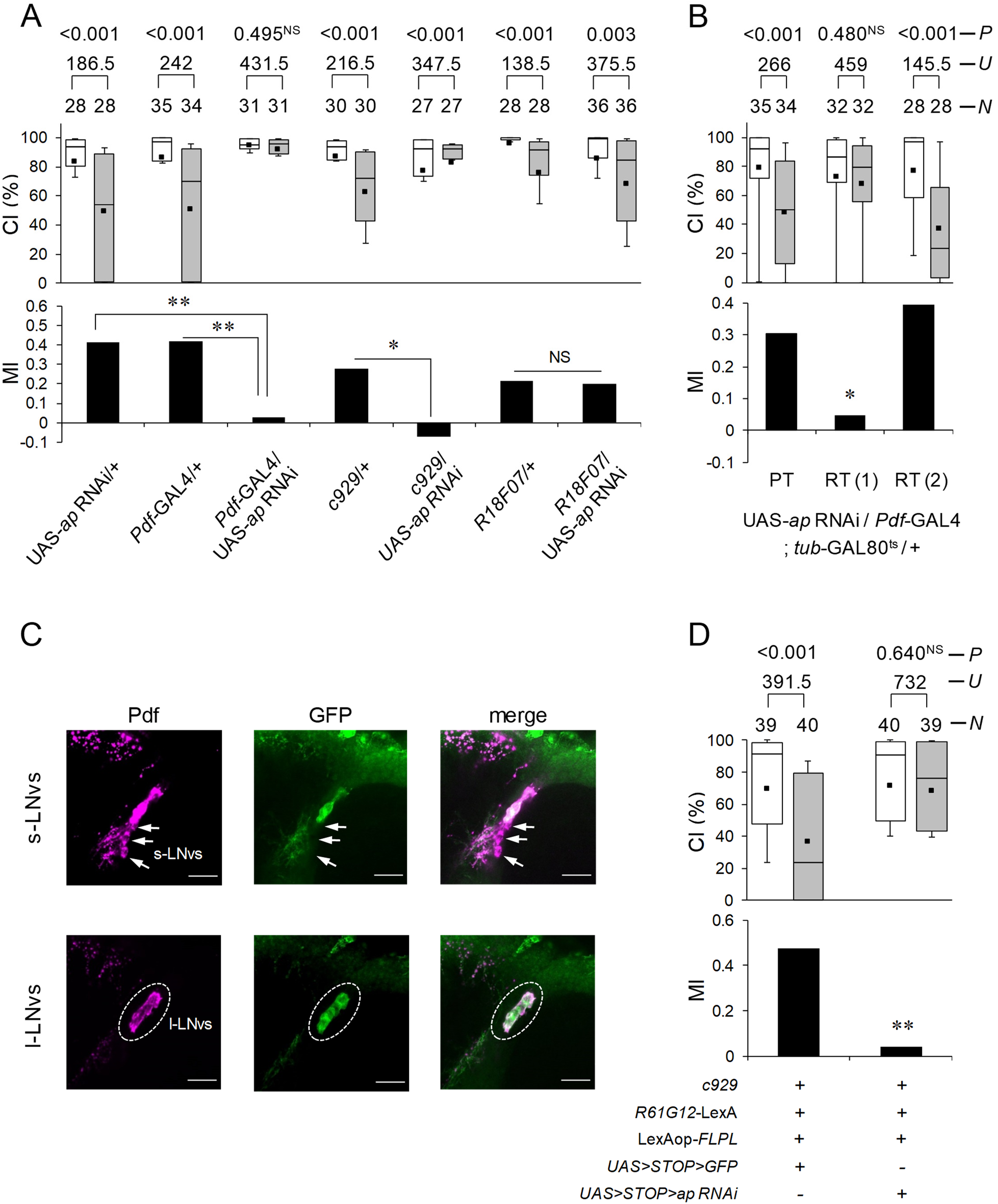
*ap* knockdown in Pdf-positive l-LNvs induces LTM impairment. ***A***, 5 d memory after 7 h conditioning. *ap* RNAi was driven by *Pdf*-GAL4, *c929*, and *R18F07*. *, *P* < 0.05; **, *P* < 0.01; NS, not significant. ***B***, 5 d memory after 7 h conditioning. UAS-*ap* RNAi/*Pdf*-GAL4; *tub*-GAL80^ts^/+ flies were used. PT, permissive temperatures (25 °C). RT (1), flies were kept at restrictive temperature (RT, 30 °C) for 24 h before the end of conditioning. RT (2), flies were kept at RT for 48–72 h after 7 h conditioning. *, *P* < 0.05; NS, not significant. ***C***, l-LNv-specific expression of mCD8::GFP. c929/*R61G12*-LexA; LexAop-*FLPL*/UAS>STOP>*mCD8::GFP* flies were used. Brains were stained with an anti-Pdf antibody and an anti-GFP antibody. Magenta, Pdf; green, mCD8::GFP. Confocal section images at the level of Pdf neurons of the adult brain. Scale bars represent 10 μm. ***D***, l-LNv-specific knockdown of *ap*. 5 d memory after 7 h conditioning. *c929*/*R61G12*-LexA; LexAop-*FLPL*/UAS>STOP>*ap* RNAi flies were used. **, *P* < 0.01; NS, not significant.

Next, we performed l-LNv-specific Ap knockdown using the Flp-out system (Rodriguez et al., 2012) with *c929* and *R61G12*-LexA. In *R61G12*-LexA, LexA is expressed in l-LNvs and s-LNvs (Inami et al., 2020). Using this system, we can knock down targeted genes specifically in the *c929* and *R61G12*-LexA-coexpressing neurons, namely, l-LNvs. The effectiveness of the Flp-out system was confirmed using the GFP reporter. In control flies carrying c929, *R61G12*-LexA, LexAop-*FLPL*, and UAS>STOP> *GFP*, the GFP signal was detected in all l-LNvs but not in s-LNvs (Fig. 4*C*). As shown in Fig. 4*D*, the Flip-out experiments with UAS>STOP>*ap* RNAi demonstrated that l-LNv-specific *ap* knockdown impaired LTM (Permutation test, *P* = 0.002), indicating that Ap in l-LNvs is indispensable for LTM.

### *ap* expression in MB α/β and Pdf neurons rescues LTM phenotype in heterozygous *ap^null^* flies

We examined whether *ap* expression in MB α/β and/or Pdf neurons rescues LTM phenotype in *ap^null^*/+ flies. First, we measured memory on day 1 after 7 h conditioning (Fig. 5*A*, 1 d memory). The targeted expression of *ap* in Pdf neurons rescued impairment of 1 day memory in *ap^null^*/+ flies (Fig. 5*A*; Permutation test; *P* < 0.001), whereas that in MB α/β neurons did not (Fig. 5*A*; Permutation test; *P* = 0.130). In contrast, Pdf neuron-specific *ap* expression did not rescue the impairment of memory on day 2 after 7 h conditioning in *ap^null^*/+ flies (Fig. 5*B*, 2 d memory; Permutation test, *P* = 0.496). These findings indicate that Pdf neuron-specific Ap expression in *ap^null^*/+ flies is not sufficient to maintain LTM for more than 2 d. For 5 d memory, the targeted expression of *ap* in either Pdf or MB α/β neurons was not sufficient to rescue the *ap^null^*/+ phenotype (Fig. 5*D*; Permutation test, vs *Pdf*-GAL4, *P* = 0.778, vs *R41C10*, *P* = 0.259). However, when *ap* was expressed in both Pdf and MB α/β neurons using *Pdf*-GAL4; *R41C10* flies (Fig. 5*C*), LTM impairment in *ap^null^*/+ flies was rescued (Fig. 5*D*; Permutation test; *P* = 0.042). These findings support the idea that Ap in Pdf neurons is responsible for memory consolidation, whereas that in MB α/β neurons is responsible for LTM maintenance.

**Figure 5.**
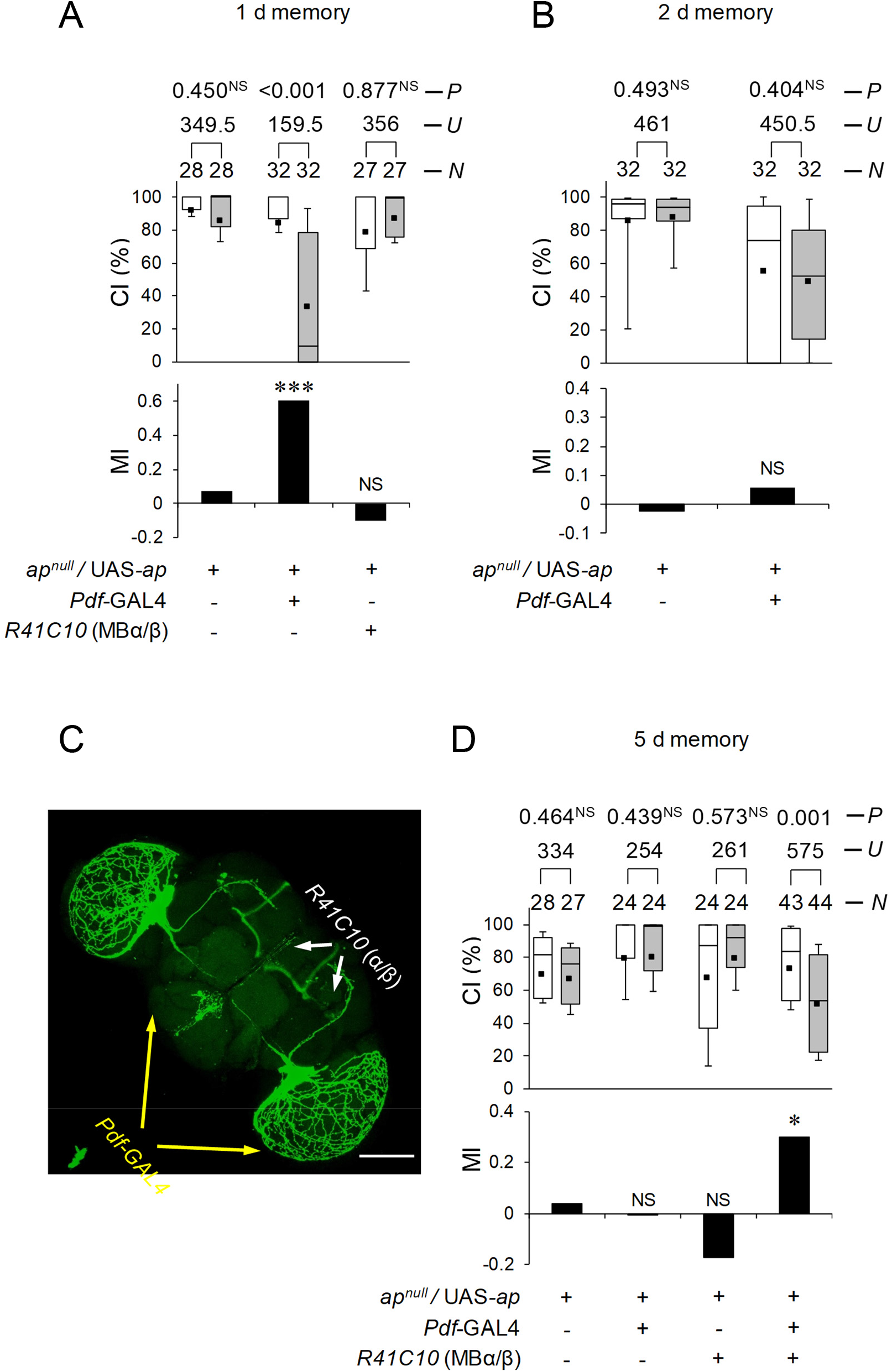
*ap* expression in MB α/β and Pdf neurons in *ap^null^*/+ flies restores LTM. ***A*, *B* and *D***, *ap^null^*/UAS-*ap* flies were used as a control. *, *P* < 0.05; ***, *P* < 0.001; NS, not significant. ***A***, 1 d memory after 7 h conditioning. ***B***, 2 d memory after 7 h conditioning. ***C***, Stacked confocal image showing an anterior view of the adult brain. A scale bar represents 100 μm. *Pdf*-GAL4/UAS-*mCD8::GFP*; *R41C10*/+ flies were used. ***D***, 5 d memory after 7 h conditioning.

### Chi in MB α/β neurons, but not in Pdf neurons, is necessary for LTM

Since the transcriptional activity of Ap requires the formation of a complex with a cofactor, Chi (van Meyel et al., 1999; van Meyel et al., 2000), it is likely that Chi is also involved in LTM. To investigate this possibility, a null allele *Chi^e5.5^* (hereafter referred to as *Chi^null^*) was used (Morcillo et al., 1997). Since *Chi^null^* is also homozygous lethal as was observed in *ap^null^*, we used heterozygous mutant flies (*Chi^null^*/+). Unlike the LTM phenotype in *ap^null^*/+, 1 d memory was not impaired in *Chi^null^*/+ flies (Fig. 6*A*; Permutation test, *P* = 0.087). Since a 50% reduction of *Chi* function in *Chi^null^*/+ flies may be insufficient to induce LTM impairment, we further knocked down *Chi* in Pdf neurons in *Chi^null^*/+ flies and examined the effect of this knockdown on LTM. The effectiveness of *Chi* RNAi was confirmed by qRT-PCR using a pan-neural GAL4 line (*nSyb*-GAL4); *Chi* RNA levels were reduced by about 55% (Fig. 6*B*; Scheffe’s multiple comparisons, GAL4 control vs F_1_, *P* < 0.001, UAS control vs F_1_, *P* < 0.001). Even when *Chi* was knocked down in a Pdf neuron-specific manner in *Chi^null^*/+ background flies, 1 d memory was not affected (Fig. 6*C*; Permutation test, *P* = 0.488). In contrast to 1 d memory, 5 d memory in *Chi^null^*/+ flies was impaired (Fig. 6*D*, Permutation test, *P* = 0.026), indicating that Chi is involved in LTM maintenance rather than memory consolidation. Furthermore, we confirmed that *Chi* knockdown in MB α/β neurons induces impairment of 5 d memory (Fig. 6*E*; Permutation test, *R41C10*/UAS-*Chi* RNAi vs UAS control, *P* = 0.007, *R41C10*/UAS-*Chi* RNAi vs GAL4 control, *P* = 0.015). Taken together, these findings support the idea that Ap cooperates with Chi in MB α/β neurons to maintain LTM, but plays a role in l-LNvs to consolidate memory independently of Chi.

**Figure 6.**
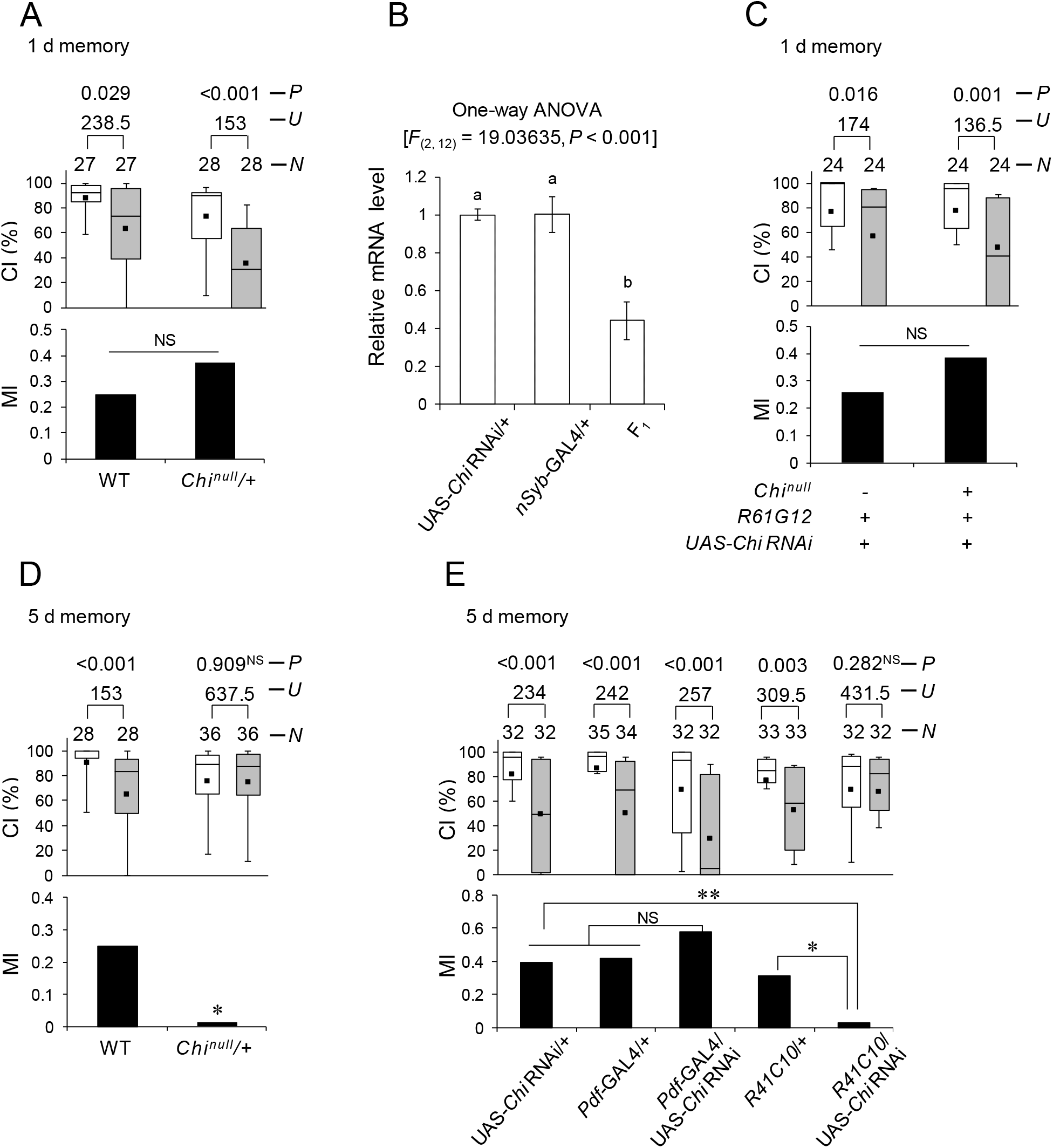
*Chi* in MB α/β is necessary for LTM. ***A***, 1 d memory after 7 h conditioning. Wild-type and *Chi^null^*/+ flies were used. NS, not significant. ***B***, Real-time qRT-PCR analysis of *ap* mRNA expression level. *nSyb*-GAL4 was used for the induction of *Chi* RNAi. Mean ± SEM was calculated from five replicates. One-way ANOVA followed by post-hoc analysis using Scheffe’s test for multiple comparisons was used. Bars with the same letter indicate values that are not significantly different (*P* > 0.05). ***C***, Pdf neuron-specific *Chi* knockdown in *Chi^null^*/+ flies. 1 d memory after 7 h conditioning. UAS-*Chi RNA/*+; *R61G12/+* (control) and UAS-*Chi RNA/Chi^null^*; *R61G12/+* flies were used. ***D***, 5 d memory after 7 h conditioning. Wild-type and *Chi^null^*/+ flies were used. *, *P* < 0.05. ***E***, *Chi* knockdown using *Pdf*-GAL4 and *R41C10*. 5 d memory after 7 h conditioning. *, *P* < 0.05; **, *P* < 0.01; NS, not significant.

### Disruption of neurotransmission and electrical silencing of Pdf neurons impairs memory consolidation

To determine whether neurotransmission from Pdf neurons is necessary for proper memory consolidation, we used the temperature-sensitive Dynamin mutation s*hibire^ts1^* (*shi^ts1^*). The targeted expression of *shi^ts1^* can inhibit neurotransmission in a spatially specific and temperature-dependent manner (Kitamoto, 2001). *Pdf*-GAL4/UAS-*shi^ts1^* flies showed LTM at the permissive temperature (PT) (Fig. 7*A*). We found that disruption of neurotransmission during the conditioning phase impaired LTM (Fig. 7*B*), but disruption during the memory maintenance or test phase did not (Fig. 7*C,D*). Thus, neurotransmission from Pdf neurons is necessary for proper memory consolidation. In addition, we performed electrical silencing of Pdf neurons using the inwardly rectifying Kir2.1 channel combined with the TARGET system. For the silencing of Pdf neurons during memory consolidation, we shifted the temperature from PT to RT and vice versa during the conditioning phase. As was observed in the disruption of neurotransmission using *shi^ts1^*, the electrical silencing of Pdf neurons also impaired LTM (Fig. 7*E*).

**Figure 7.**
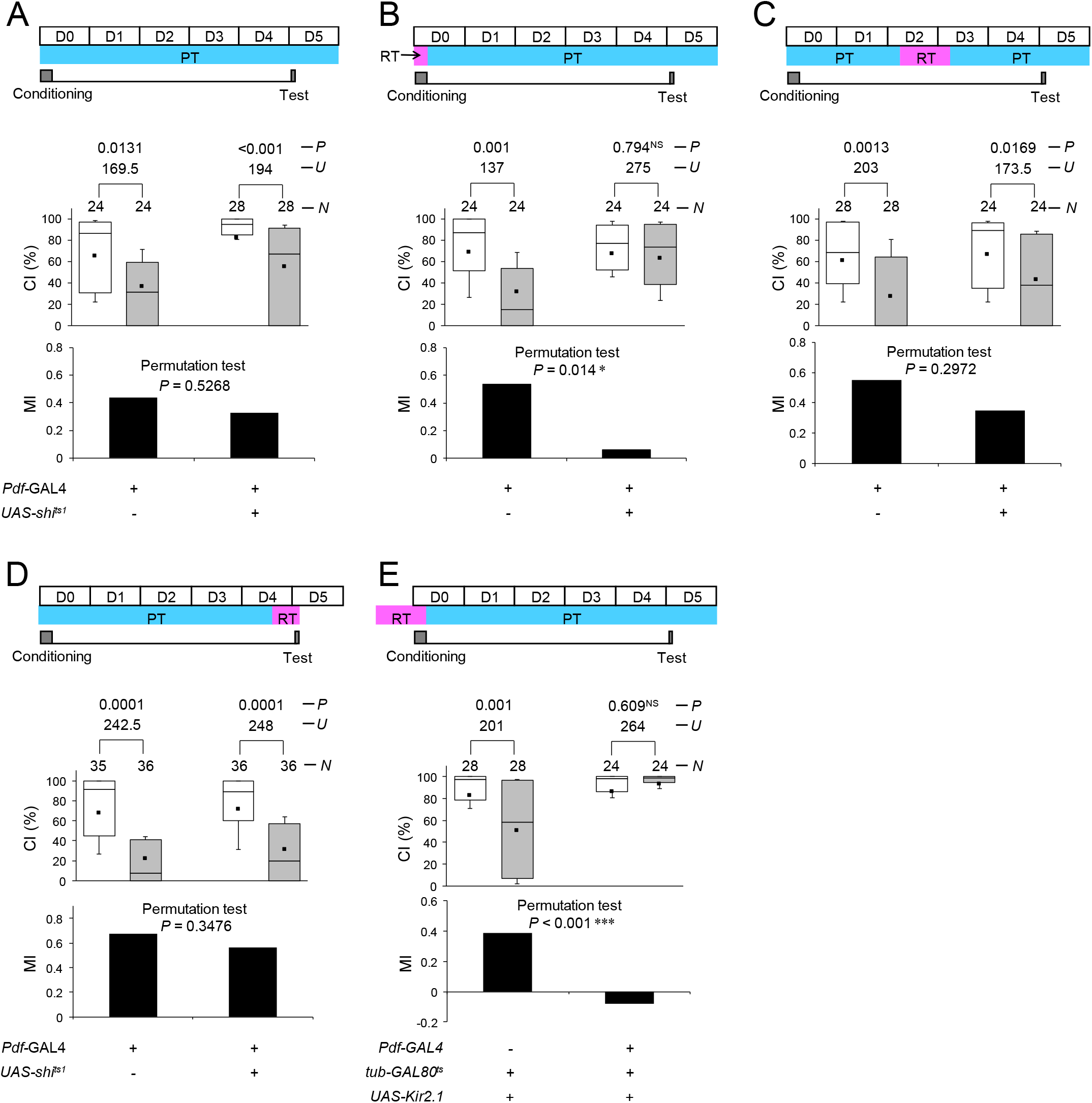
Disruption of neurotransmission and electrical silencing of Pdf neurons impairs memory consolidation. ***A–D***, 5 d memory after 7 h conditioning. *Pdf*-GAL4/+ (control) and *Pdf*-GAL4/UAS-*shi^ts1^* flies were used. *, *P* < 0.05; ***, *P* < 0.001; NS, not significant. ***A***, All experiments were performed at PT (25 °C). ***B***, Flies were kept at RT during conditioning. ***C***, Flies were kept at RT for 48–72 h after 7 h conditioning. ***D***, Flies were kept at RT for 12 h before the test. ***E***, 5 d memory after 7 h conditioning. Flies were kept at RT for 48–72 h after 7 h conditioning. UAS-*Kir2.1*/*Pdf*-GAL4; *tub*-GAL80^ts^/+ flies were used. UAS-*Kir2.1*/*tub*-GAL80^ts^ flies were used as the control.

### Ap in Pdf neurons is indispensable for appropriate response to GABA

Our results show that Ap in Pdf neurons is necessary for memory consolidation in a Chi-independent manner (Fig. 3, 5, and 6). Since all temporal genetic manipulations in Pdf neurons (Ap knockdown, disruption of neurotransmission, and electrical silencing) impair memory consolidation, *ap* mutations or knockdown may reduce the excitability of Pdf neurons. A *Drosophila* ionotropic GABA_A_ receptor (GABA_A_R), Resistant to Dieldrin (Rdl), is expressed in Pdf neurons, and anti-RDL antibody staining reveals that RDL is highly expressed in l-LNv somata, while little or no expression is observed in s-LNv somata (Parisky et al., 2008; Chung et al., 2009). Furthermore, the electrophysiological analysis revealed that l-LNvs respond to GABA (Chung et al., 2009). Considering the physiological properties of l-LNvs, we next sought to investigate the possibility that Ap in l-LNvs is involved in response to GABA.

To visualize the response to GABA in l-LNvs, the FRET-based Cl^−^ probe, SuperClomeleon, was used (Grimley et al., 2013). First, in flies with wild-type *ap*, we confirmed robust Cl^−^ responses to 400 μM GABA in l-LNvs in the presence of TTX, while the responses are blocked by the GABA_A_R antagonist picrotoxin (Fig. 8*A, B*). Similar Cl^−^ responses were observed in heterozygous *Chi^null^* flies (Fig. 8*C*, *D*). Compared with wild-type and heterozygous *Chi^null^* flies, heterozygous *ap^null^* flies showed robust increases in Cl^−^ responses (Fig. 8*C, D*). These results indicate that Ap is necessary for proper responses to GABA in l-LNvs. If the reduced Ap expression in *ap^null^*/+ decreases LNv excitability due to overresponse to the inhibitory neurotransmitter GABA and the inhibition of LNv-excitability causes the 1 d memory impairment, 1 d memory phenotype in heterozygous *ap^null^* flies may be compensated by reduced *Rdl* expression in l-LNvs. As we expected, the knockdown of *Rdl* in Pdf neurons compensated the impairment of 1 d memory in *ap^null^*/+ flies (Fig. 8*E*). Furthermore, the Pdf neuron-specific overexpression of *Rdl* in wild-type background induced 1 d memory impairment (Fig. 8*F*). This result is consistent with the aversive effect of the electrical silencing of Pdf neurons on LTM.

**Figure 8.**
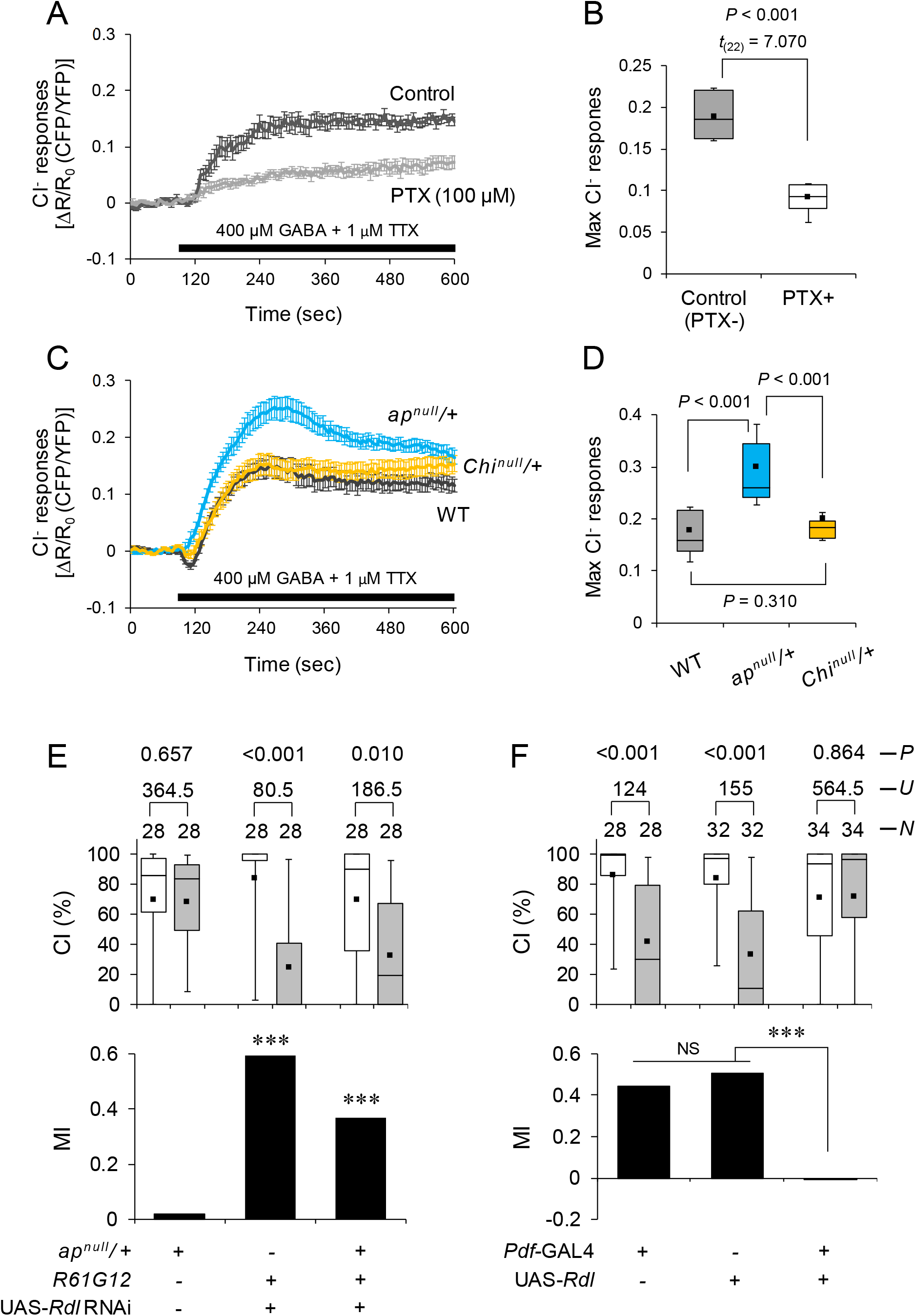
Ap in l-LNvs is indispensable for appropriate response to GABA. ***A–D***, *Ex vivo* imaging using SuperClomeleon. ***A, B***, UAS-SuperClomeleon/*R61G12* flies were used. PTX, picrotoxin. ***A***, Traces of mean SuperClomeleon response to 400 μM GABA and 1 μM TTX with or without bath-applied 100 μM PTX. Error bars, SEM; *N* = 12 in each trace. ***B***, Maximum percentage change in fluorescence of the SuperClomeleon related to ***A***. Student’s *t*-test was used. ***C***, UAS-SuperClomeleon/*R61G12* flies (WT), UAS-SuperClomeleon/*ap^null^*; *R61G12*/+ (*ap^nill^*/+), and UAS-SuperClomeleon/*Chi^null^*; *R61G12*/+ (*Chi^null^*/+) flies were used. Traces of mean SuperClomeleon response to 400 μM GABA and 1 μM TTX. Error bars, SEM; *N* = 20-24 in each trace. ***D***, Maximum percentage change in fluorescence of the SuperClomeleon related to ***C***. Nonparametric ANOVA (Kruskal–Wallis test) followed by post-hoc analysis using the Steel–Dwass test was carried out for multiple comparisons. ***E* and *F***, 1 d memory after 7 h conditioning. ***, *P* < 0.001; NS, not significant. ***E***, *Rdl* knockdown in Pdf neurons in *ap^null^*/+. *ap^null^*/+, UAS-*Rdl* RNAi/*R61G12*, and *ap^null^*/UAS-*Rdl* RNAi; *R61G12* flies were used. ***F***, *Rdl* overexpression in Pdf neurons. *Pdf*-GAL4/+, UAS-*Rdl/+*, and *Pdf*-GAL4/UAS-*Rdl* flies were used.

## Discussion

The evolutionary conserved LIM-HD protein Ap acts as a transcriptional activator, and it is essential for various developmental events in *Drosophila* (Cohen et al., 1992; Diazbenjumea and Cohen, 1993; Lundgren et al., 1995; van Meyel et al., 2000). However, the functions of centrally expressed Ap have remained largely unknown. In this study, we identified novel Ap functions. First, Ap and its cofactor Chi in the MB α/β neurons are indispensable for LTM maintenance. Ap LIM domains interact with the Chi LIM interaction domain, and Chi can homodimerize through the dimerization domain (van Meyel et al., 1999; Hobert and Westphal, 2000; van Meyel et al., 2000) (Fig. 9A). Since Ap/Chi regulates the transcription of Ap target genes (Hobert and Westphal, 2000), it is most likely that Ap/Chi in MB α/β neurons is necessary to provide proteins required for the maintenance of courtship LTM (Fig. 9*A*). Second, unlike the role of Ap in MB α/β neurons, Ap in Pdf-positive l-LNvs was essential for memory consolidation to establish courtship LTM in a Chi-independent manner. In *Drosophila* wing development, it is proposed that Chi binds to Ap and that Beadex (Bx) [also known as *Drosophila* LIM-only protein (dLMO)] modulates Ap function by interfering with the formation of Ap/Chi (Milan et al., 1998). In this scenario, Bx can modify Ap-dependent transcription in the presence of Chi. In the adult fly brain, *Bx* expression is prominent in several distinct brain regions including the MBs and Pdf neurons (Heberlein et al., 2009). Since the Ap/Chi complex is critical for LTM maintenance in the MB α/β neurons, Bx may be involved in LTM maintenance by regulating the activity of the Ap/Chi complex. In fact, in aversive olfactory memory, Bx in MB neurons is involved in LTM maintenance (Hirano et al., 2016). Thus, Bx may also affect the maintenance of courtship LTM via modification of Ap/Chi-dependent transcription in MB α/β neurons. Interestingly, an Ap function newly identified in l-LNvs is independent of Chi. Thus, the molecular mechanisms responsible for the Ap function in l-LNvs should be different from those in the previously reported model. Further studies are required to determine how memory consolidation is regulated by the Chi-independent Ap function in Pdf neurons.

**Figure 9.**
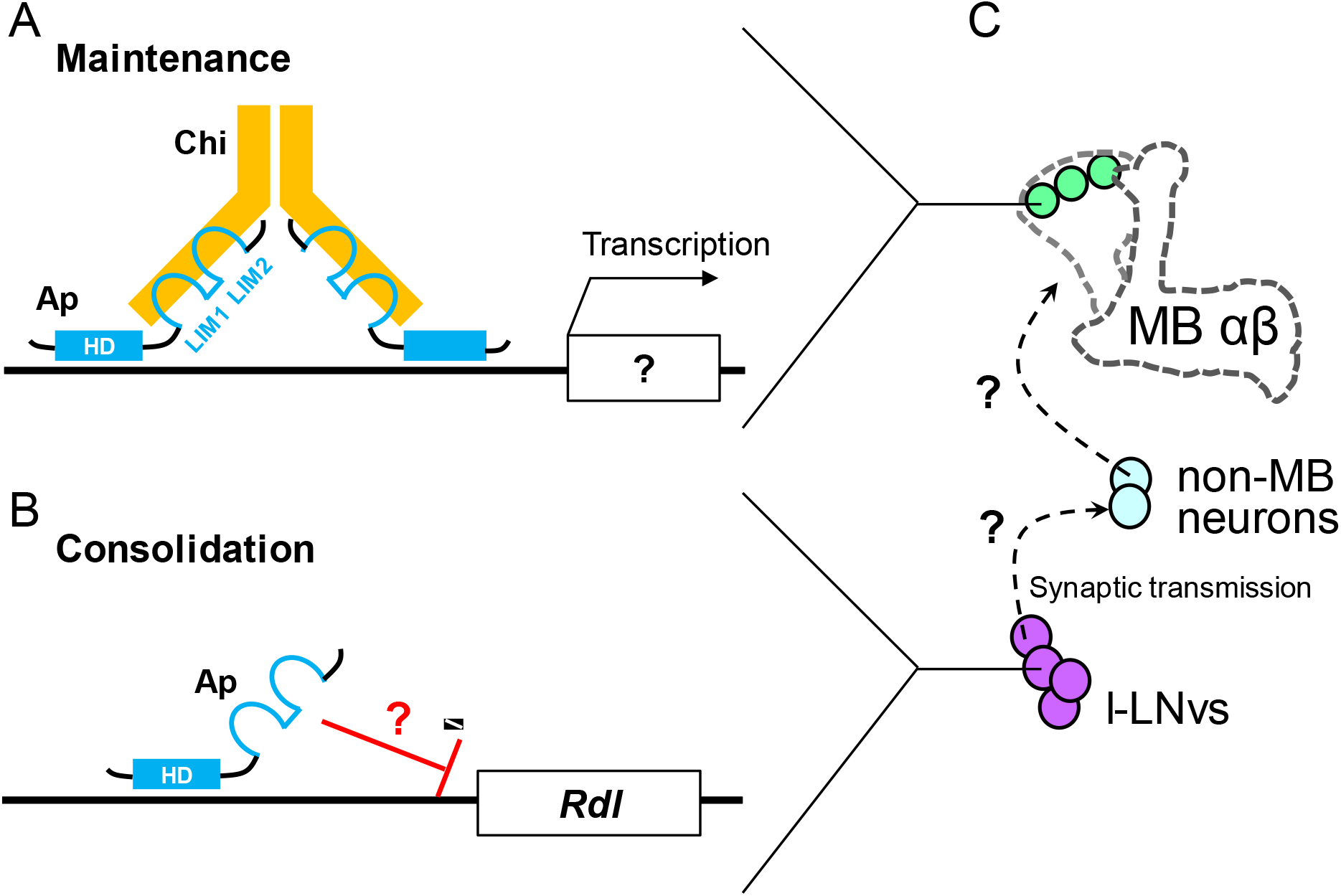
Possible model of Ap functions in the consolidation and maintenance of LTM. ***A***, Ap/Chi complex induces gene expression. Subsequently, proteins required for maintaining consolidated LTM are produced in MB α/β neurons. ***B***, In l-LNvs, Ap may inhibit *Rdl* expression in a Chi-independent manner. ***C***, Intercellular communication from l-LNvs to MB neurons plays a crucial role in forming a stable LTM. Non-MB neurons may mediate intercellular communication between l-LNvs and MB neurons because l-LNvs do not project to MBs directly.

As for Chi-independent Ap function, we provide evidence that Ap is necessary for appropriate Cl^−^ responses to GABA in l-LNvs; *ap* null mutation enhanced the Cl^−^ responses in l-LNvs. Consistent with the finding that the Ap function is independent of Chi in l-LNvs, *Chi* null mutation did not affect the Cl^−^ response to GABA. Considering the Ap mutant phenotype, wild-type Ap likely contributes to reducing l-LNv responses to GABA and maintains higher excitability of l-LNvs. We also confirmed that *Rdl* knockdown in Pdf neurons compensates for the defective 1 d memory observed in heterozygous *ap* null mutant flies. These findings suggest that Ap in the wild-type reduces Cl^−^ influx by suppressing the expression or modulating the channel properties of GABA_A_R in l-LNvs. Some LIM-HD proteins (e.g., Lhx2, Lhx6, and Islet-1) acts as transcriptional repressors as well as transcriptional activators (Gay et al., 2000; Subramanian et al., 2011; Zhang et al., 2013). Thus, it will be interesting to determine whether Ap also acts as a transcriptional repressor and inhibits the transcription of *Rdl* encoding the *Drosophila* GABA_A_R in l-LNvs (Fig. 9*B*).

As is well known in the olfactory associative learning paradigm, MB neurons are critically involved in memory consolidation to establish courtship LTM (Ishimoto et al., 2009; Sakai et al., 2012; Fitzsimons et al., 2013; Kruttner et al., 2015; Inami et al., 2020). In this study, we identified that synaptic transmission from l-LNvs is necessary for memory consolidation (Fig. 7), suggesting that intercellular communication from l-LNvs to MB neurons plays a crucial role in establishing a stable LTM. The l-LNvs arborize in the most distal layer of the medulla and the accessory medullae (aMe), and they project to the other hemisphere through the posterior optic tract (POT) (Helfrich-Forster et al., 2007). Thus, it is unlikely that l-LNvs synaptically project and transmit to the MB α/β neurons directly (Fig. 9*C*).

Sleep is essential for the consolidation of *Drosophila* courtship memory (Ganguly-Fitzgerald et al., 2006; Donlea et al., 2011). In addition, Pdf-positive l-LNvs regulate light-dependent arousal (Shang et al., 2008), and knockdown of Ap and Chi in l-LNvs reduces sleep amount (Shimada et al., 2016). An interesting question now arises: Does reduced sleep caused by Ap knockdown impair memory consolidation? Unlike *ap* knockdown, *Chi* knockdown in Pdf neurons did not affect 1 d memory (Fig. 6), even though it reduces the sleep amount (Shimada et al., 2016). Thus, impairment of memory consolidation induced by Ap knockdown in l-LNvs does not merely result from reduced sleep.

We previously identified that the targeted expression of a CREB repressor in MB α/β neurons during the memory maintenance phase (48–72 h after conditioning) impairs LTM (Inami et al., 2020), indicating that CREB-dependent transcription in MB α/β neurons is necessary for the maintenance of courtship LTM. Similarly, in this study, our results support the idea that Ap/Chi-dependent transcription in MB α/β neurons contributes to the maintenance of LTM for more than 2 d. Taken together, courtship LTM is likely stored in MB α/β neurons from the second day after conditioning, and proteins required for mainaining LTM for more than 2 d should be provided via transcriptions by CREB and Ap/Chi. Although molecular interactions between CREB and Ap/Chi remain unclarified, it will be of interest to investigate whether the transcription by CREB and that by Ap/Chi occur in MB α/β neurons sequentially or in parallel.

The Pdf neuropeptide is necessary for the maintenance of courtship LTM, and electrical silencing of Pdf neurons using Kir2.1 impairs the LTM maintenance (Inami et al., 2020). However, we found that disruption of the Dynamin function using *shi^ts1^* in Pdf neurons impairs memory consolidation but not LTM maintenance (Fig. 7). These seemingly paradoxical consequences may result from the characteristic difference between Kir2.1 and mutated Dynamin (Shi^ts1^) in neurotransmission. Since the induction of *shi^ts1^* in neurons blocks synaptic vesicle recycling at the restrictive temperature, neurotransmission in target neurons should be disrupted (Kitamoto, 2001). The induction of *Kir2.1* in target neurons suppresses the excitability of the target neurons. Consequently, neurotransmission from the target neurons should be blocked (Hodge, 2009). Although the Pdf neuropeptide is essential for the generation of locomotor activity rhythms in *Drosophila* (Renn et al., 1999), disruption of the Dynamin function in Pdf neurons has little effect on locomotor activity rhythms (Mabuchi et al., 2016). However, the temporal induction of Kir2.1 in Pdf neurons can induce arrhythmic locomotor activity (Depetris-Chauvin et al., 2011). On the basis of these findings, it is considered that Kir2.1 blocks Pdf release, but Dynamin-dependent synaptic transmission has little effect on Pdf release. We previously identified by qRT-PCR analysis that Pdf neuron-specific Ap knockdown does not affect Pdf expression (Shimada et al., 2016). Furthermore, Pdf knockdown impairs LTM maintenance; nevertheless it does not affect memory consolidation (Inami et al., 2020). Considering these findings, it is unlikely that Pdf release from l-LNvs is involved in memory consolidation. Our findings raise the possibility that Dynamin-dependent synaptic transmission (e.g., classical neurotransmitter release) from l-LNvs contributes to memory consolidation for the establishment of courtship LTM (Fig. 9*C*).

In *Drosophila*, the LTM maintenance phase has been conceptually defined as the time after LTM is fully formed and consolidated, and it is generally believed that memory consolidation is completed within 1 d after conditioning (Margulies et al., 2005; Davis, 2011). Hirano et al. have reported that the consolidation and maintenance phases are molecularly separated in aversive olfactory LTM (Hirano et al., 2016). Similarly, we have proposed that the maintenance phase in courtship LTM is also molecularly distinguished from the memory consolidation phase (Inami et al., 2020). Because *Pdf^01^* mutant flies display intact 1 d memory but are defective in 2 d memory (Inami et al., 2020), it is possible that the maintenance phase can be distinguished from the consolidation phase on the basis of the requirement of Pdf function. In this study, we found that *ap^null^*/*Pdf*-GAL4; UAS-*ap*/+ flies also show intact 1 d memory but 2 d memory impairment (Fig. 5), suggesting that the consolidation and maintenance phases in courtship LTM are molecularly and cellularly separated.

Ap is evolutionally conserved in vertebrates and invertebrates, and its function in regulating developmental events is also well conserved (Hobert and Westphal, 2000). In this study, we found that consolidation and maintenance of LTM require distinct Ap functions in fly clock neurons and the memory center. Although little is known about the functional significance of mammalian Ap orthologues in the adult brain, our findings raise the interesting possibility that they are also involved in multiple phases of memory processes.

## Materials and Methods

### Fly stocks

All flies were raised on glucose–yeast–cornmeal medium in 12:12 LD cycles at 25.0 ± 0.5 °C (45–60% relative humidity). Virgin males and females were collected without anesthesia within 8 h after eclosion. The fly stocks used for this study were as follows: wild-type Canton-S (CS), *ap^rk568^* (BL5374), *ap^UGO35^* (provided by Dr. Manfred Frasch, University of Erlangen-Nuremberg), *ap::GFP* (BL38423), *Chi^e5.5^* (BL4541), *Pdf*-GAL4 (BL6900), *c929* (BL25373), *R18F07* (BL47876), *R61G12* (BL41286), *R61G12*-LexA (BL52685), *R41C10* (BL50121), *R55D03* (BL47656), *c305a* (BL30829), *nSyb*-GAL4 (BL51941), hs-GAL4 (Sakai et al., 2012), UAS-*shi^ts1^* (Kitamoto, 2001), UAS-*Kir2.1::eGFP* (BL6596), UAS-*mCD8::GFP* (BL5137), UAS-*mCherry::NLS* (BL38424), UAS-*ap* RNAi (NIG-fly, 8376R-1), UAS-*ap* (see next section), UAS-*Chi* RNAi (VDRC, 43934), UAS-*Rdl* (BL29036), UAS-*Rdl* RNAi (VDRC, 41103), UAS-*SuperClomeleon* (BL59846), *tub*-GAL80^ts^ (BL7017), LexAop2-*FLPL* (BL55820), UAS>stop>*mCD8::GFP* (BL30032), and UAS>stop>*ap* RNAi (see next section). All lines used for behavior experiments except for *ap^UGO35^*, *Chi^e5.5^*, and UAS>stop>*mCD8::GFP* were outcrossed for at least five generations to *white^1118^* flies with the CS genetic background.

### Generation of transgenic flies

Full-length *ap* cDNA was isolated by RT-PCR with adult fly head RNA and two primers, 5’-GCGGCCGCCAAAATGGGCGTCTGCACCGAGGAGCGC-3’ and 5’-TCTAGATTAGTCCAAGTTAAGTGGCGGTGTGC-3’. The PCR product was digested with NotI and XbaI, and cloned into a pBluescript (pBS) II SK(+). The *ap* cDNA was subcloned into the NotI/XbaI-digested pUAST attB vector (Bischof et al., 2007). Constructs of UAS-*ap* were injected into eggs of PBac{y[+]-attP-9A}VK00005 (BL24862).

For the generation of the UAS-FRT-stop-FRT*-ap* RNAi construct (UAS>stop>*ap* RNAi), an *ap* RANi fragment was obtained by PCR from the genomic DNA of UAS-*ap* RNAi (NIG-fly, 8376R-1). Subsequently, it was inserted into the pUAST attB vector using an In-Fusion^®^ HD Cloning Kit (Clontech). The primer sequences used for the PCR are as follows: forward, 5’-AACAGATCTGCGGCCGCATAACGCGCAACCTCGAC-3’; reverse, 5’-ACAAAGATCCTCTAGAGGAACAATGCTCCGACTAG-3’. Next, the FRT-stop-FRT cassette (<stop<) was PCR-amplified from the UAS<stop<*mCD8::GFP* vector (Addgene, 24385) and then cloned into the pUAST attB vector containing the *ap* RNAi fragment by using an In-Fusion^®^ HD Cloning Kit. The primer sequences used for the PCR are as follows: forward, 5’-AGGGAATTGGGAATTCGAAGTTCCTATTCCGAAG-3’; reverse, 5’-ATCTGTTAACGAATTCGAAGTTCCTATACTTTCTAG-3’. The UAS>stop>*ap* RNAi construct was also injected into eggs of PBac{y[+]-attP-9A}VK00005.

### Courtship conditioning assay

The courtship conditioning assay was carried out as previously described (Inami et al., 2020). The male courtship activity of individual flies was calculated manually as a courtship index (CI), defined as the percentage of time spent in performing courtship behaviors for 10 min. We first measured CI of conditioned and naïve males (CI_Conditioned_ and CI_Naïve_), and then mean CI_Naïve_ was calculated. To quantify courtship memory, memory index (MI) was calculated using the following formula: MI = (mean CI_Naïve_ - mean CI_Conditioned_) / mean CI_Naïve_.

### Heat-shock treatment

In F_1_ males between hs-GAL4 and UAS-*ap* RNAi flies, the *ap* expression was inhibited after heat-shock treatment. For qRT-PCR analysis, hs-GAL4/+, UAS-*ap* RNAi/+, and hs-GAL4/ UAS-*ap* RNAi flies were heat-shocked (37 °C) for 20 min, and total RNA was extracted 3 h after heat-shock treatment. For courtship conditioning assay, hs-GAL4/ UAS-*ap* RNAi flies were heat-shocked 3 h before conditioning, immediately after conditioning, or 3 h before the test.

### Temporal disruption of neurotransmission

The temperature-sensitive allele *shibire^ts1^* (*shi^ts1^*) is defective in Dynamin function at restrictive temperatures (RT). In F_1_ males between *Pdf*-GAL4 and UAS-*shi^ts1^*, *shi^ts1^* is expressed in Pdf neurons. In these males, neurotransmission in Pdf neurons should be disrupted at RT but not at permissive temperatures (PT, 25 °C). To disrupt neurotransmission in Pdf neurons during the memory consolidation, maintenance, or test phase, the temperature was increased to RT (32 °C) during the three experimental periods: conditioning phase, 48–72 h after conditioning (memory maintenance phase), and 12 h before the test initiation.

### Temporal gene expression using the TARGET system

The *tub*-GAL80^ts^ transgene used in the TARGET system (McGuire et al., 2003) encodes a ubiquitously expressed and temperature-sensitive GAL4 repressor that is active at PT (25 °C) but not at RT (30 °C). By using UAS-*ap* RNAi combined with the TARGET system, we knocked down *ap* in GAL4-positive neurons at RT, but not at PT. We shifted PT to RT and vice versa during two experimental phases: 24 h before the end of conditioning and 48–72 h after conditioning. To drive the expression of *Kir2.1* in Pdf neurons during memory consolidation, flies were kept at RT during conditioning.

### *ap* knockdown in large ventral lateral clock neurons

Pdf neurons are classified into two neural clusters, s-LNvs and l-LNvs. To assay whether l-LNv-specific *ap* knockdown affects LTM, two binary gene expression systems (GAL4/UAS and LexA/LexAop) combined with Flippase (FLP/FRT) were used. The specific target gene is expressed in GAL4- and LexA-coexpressing neurons utilizing this system. *R61G12*-LexA, *c929*, and UAS>stop>*ap* RNAi lines were used in the experiments.

### Real-time quantitative reverse transcription PCR (qRT-PCR)

For *ap*, TRizol (Invitrogen) was used for collecting total RNA from about 30 male fly heads in each genotype. For *Chi*, a PicoPure RNA Isolation Kit (KIT0204, Thermo Fisher Scientific) was used for collecting total RNA from three whole brains in each genotype. cDNA was synthesized by the reverse transcription reaction using a QuantiTect Reverse Transcription Kit (#205311, QIAGEN). qRT-PCR was carried out using a Chromo 4 detector (CFB-3240, MJ Research) and the SYBR Premix Ex Taq™ (Takara Bio Inc.) for *ap* or the THUNDERBIRD SYBR qPCR Mix (QPS-201, TOYOBO) for *Chi*. The primer sequences (custom-made by Eurofins Genomics) used for qRT-PCR were as follows: *ap*-Forward, 5’-ATAACGCGCAACCTCGACGAC-3’; *ap*-Reverse, 5’-CATGAGGATTCCCGTTCCAGC-3’; *Chi*-Forward, 5’-AACGGGCCGTGAAAAGTGTG-3’; *Chi*-Reverse, 5’-GTGGTCGGTTCTATCGGGCA-3’; *rp49*-Forward, 5’-AAGATCGTGAAGAAGCGCAC-3’; *rp49*-Reverse, 5’-TGTGCACCAGGAACTTCTTG-3’. The expression level of each mRNA was normalized to that of *rp49* mRNA. The average of the normalized mRNA expression levels in control flies was calculated using data from 5 or 6 independent assays.

### Immunohistochemistry

Immunohistochemistry was performed as previously described (Shimada et al., 2016). For Pdf staining, brains were stained with a mouse anti-Pdf antibody (PDF C7-s, Developmental Studies Hybridoma Bank at the University of Iowa, 1:200) followed by Alexa Fluor 568 anti-mouse IgG (A11004, Thermo Fisher Scientific) as the secondary antibody (1:1000). For GFP staining, brains were stained with a rabbit anti-GFP antibody (A11122, Thermo Fisher Scientific, 1:200), followed by Alexa Fluor 488 anti-rabbit IgG (A11008, Thermo Fisher Scientific, 1:1000) as the secondary antibody. Fluorescence signals were observed under a confocal microscope [(LSM710, Zeiss) or (C2, Nikon Corp.)].

### Functional fluorescence imaging

Brains were prepared for imaging analysis as previously described with some modifications (Sato et al., 2018). Briefly, brains were extracted in ice-cold Ca^2+^-free HL3 medium (in mM, NaCl, 70; sucrose, 115; KCl, 5; MgCl_2_, 20; NaHCO_3_, 10; trehalose, 5; Hepes, 5) (Stewart et al., 1994). Phenol red was used as a pH indicator (final concentration, 0.5 mg/ml) to adjust the pH of buffers by coloring. For adjustment of pH, CO_2_ gas was dissolved in buffers so that the color became orange (pH 7.0 ± 0.2). The brains were treated with papain (10 U/ml, activated by 15 min incubation with 0.8 mM EDTA at 37 °C) for 15 min at room temperature and then washed several times with Ca^2+^-free HL3 medium. After papain treatment, the brains were incubated with standard *Drosophila* HL3 medium (in mM, CaCl_2_, 1.8; NaCl, 70; sucrose, 115; KCl, 5; MgCl_2_, 20; NaHCO_3_, 10; trehalose, 5; Hepes, 5) with TTX alone (1 μM) or with TTX (1 μM) and PTX (100 μM). During incubation, fresh medium was infused into the chamber using a peristaltic pump (AC-2110, ATTO Corporation) every 5–10 min.

For Cl^−^ imaging using SuperClomeleon, fluorescence images were captured at 0.5 Hz at 5 s intervals using a confocal microscope system (C2, Nikon Corp.) equipped with a 20× dry objective (NA, 0.75; Nikon Corp.). The cyan fluorescent protein (CFP) was excited at 458 nm and detected using a 482 ± 17.5 nm band-pass filter, and the yellow fluorescent protein (YFP) was detected simultaneously using a 540 ± 15 nm band-pass filter. GABA was diluted in the incubation medium so that the final concentration became 400 μM, and 1.6 ml of the GABA solution was perfused 90 s after the start of capturing images. For the calculation of fluorescence resonance energy transfer (FRET) changes, the following analysis was performed using custom-made software developed in MATLAB (MathWorks). Regions of interest (ROIs) were selected as a circle within or surrounding each l-LNv soma to measure the average F of CFP and YFP at each time point. We calculated FRET changes as R using the following formula: R = average F of CFP/average F of YFP. We calculated the initial R (R_0_) by averaging the R values recorded from 10 sequential frames before stimulation. To obtain ΔR/R_0_ (%) as Cl^−^ responses, we calculated (R – R_0_)/R_0_ × 100 at each time point.

### Statistical analyses

All the statistical analyses were performed using IBM SPSS Statistics 22 (IBM Japan, Ltd.) or BellCurve for Excel (Social Survey Research Information Co., Ltd.), except for the comparisons of MI. In all statistical analyses except for the comparisons of MI, the Kolmogorov–Smirnov test was used to determine whether the data are typically distributed. In the statistical analysis of CI, when basic data were not distributed normally, we carried out the log transformation of the data. Even though the basic data or transformed data are normally distributed, heteroscedasticity was detected in all data. Thus, we used the Mann–Whitney *U* test for comparisons. In the statistical analysis of MI, the permutation test with 10000 random permutations was used (H_0_, the difference between experimental and control groups is 0). The free statistical package R was used for these tests (Koemans et al., 2017). In qRT-PCR of *ap* knockdown, Student’s *t*-test or the Mann–Whitney *U* test was used for comparisons between HS- and HS+. In qRT-PCR of *Chi* knockdown, one-way ANOVA followed by post-hoc analysis using Scheffe’s test was used for multiple comparisons. In functional fluorescence imaging, the basic data related to picrotoxin treatment were distributed normally, and homoscedasticity was evident. Thus, Student’s *t*-test was carried out. In multiple comparisons among three genotypes (wild-type, *ap^nill^*/+, and *Chi^null^*/+ flies), heteroscedasticity was evident. Thus, nonparametric ANOVA (Kruskal–Wallis test) followed by post-hoc analysis using the Steel-Dwass test was carried out.

## Acknowledgments

We thank Yuto Kurata and Keiko Irisawa for technical assistance. We also thank Toshiro Aigaki for helpful discussions. This work was supported by JSPS KAKENHI Grants (Numbers 18H04887 and 16H04816 to T.S. and 15J06303 to S.I.).

## Author contributions

Show Inami, Formal analysis, Investigation, Funding acquisition, Visualization, Methodology, Writing— review and editing; Tomohito Sato, Formal analysis, Investigation, Visualization, Methodology, Writing— review and editing; Yuki Suzuki, Formal analysis, Investigation, Methodology, Writing—review and editing; Toshihiro Kitamoto, Conceptualization, Supervision, Writing—original draft, Writing—review and editing; Takaomi Sakai, Conceptualization, Formal analysis, Supervision, Funding acquisition, Investigation, Methodology, Writing—original draft, Project administration, Writing—review and editing

